# Identification of β2 microglobulin, the product of B2M gene, as a Host Factor for Vaccinia Virus Infection by Genome-Wide CRISPR genetic screens

**DOI:** 10.1101/2022.08.10.503559

**Authors:** Alejandro Matía, Maria M. Lorenzo, Yolimar C. Romero-Estremera, Juana M. Sanchez-Puig, Angel Zaballos, Rafael Blasco

**Affiliations:** Departamento de Biotecnología, Instituto Nacional de Investigación y Tecnología Agraria y Alimentaria – Consejo Superior de Investigaciones Científicas (INIA – CSIC), Madrid, Spain; Unidad de Genómica, Centro Nacional de Microbiología-ISCIII, Madrid, Spain

## Abstract

Genome-wide genetic screens are powerful tools to identify genes that act as host factors of viruses. We have applied this technique to the analyze the infection of HeLa cells by Vaccinia virus, in an attempt to find genes necessary for infection. Infection of cell populations harboring single gene inactivations resulted in no surviving cells, suggesting that no single gene knock-out was able to provide complete resistance to Vaccinia virus and thus allow cells to survive infection. In the absence of an absolute infection blockage, we explored if some gene inactivations could provide partial protection leading to a reduced probability of infection. Multiple experiments using modified screening procedures involving replication restricted viruses led to the identification of multiple genes whose inactivation potentially increase resistance to infection and therefore cell survival. As expected, significant gene hits were related to proteins known to act in virus entry, such as ITGB1 and AXL as well as genes belonging to their downstream related pathways. Additionally, we consistently found β_2_-microglobulin, encoded by the B2M gene, among the screening top hits, a novel finding that was further explored. Inactivation of B2M resulted in 54% and 91% reduced VV infection efficiency in HeLa and HAP1 cell lines respectively. In the absence of B2M, while virus binding to the cells was unaffected, virus internalization and early gene expression were significantly diminished. These results point to β_2_-microglobulin as a relevant factor in the Vaccinia virus entry process.

**Author summary:** Orthopoxviruses, a genus belonging to the family *Poxviridae*, include human pathogens like Variola virus, the causative agent of the now eradicated Smallpox, and Monkeypox virus that cause human outbreaks of zoonotic origin. Being the prototype Poxvirus, *Vaccinia virus* has been extensively used as the ideal model to study infection. For Poxviruses, both fluid phase endocytosis and direct fusion at the plasma membrane have been described as modes of entry. To date, only a few cellular factors have been identified in the vaccinia virus entry pathway. In this study, we report that blind genome-wide genetic screens allowed us to identify several cellular factors involved in Vaccinia Virus infection, of which many could be related to known factors in virus entry. In addition, we found that β_2_-microglobulin constitute a novel player for Poxvirus entry not related to previously described cellular pathways involved in the entry process. These findings add new information to the complex picture of Poxvirus entry and open the door to the discovery of new entry mechanisms used by Poxviruses.

## Introduction

Poxviruses are dsDNA virus whose replication cycle occur entirely in the cytoplasm of the host cell (1, 2). Among the Poxviruses, Vaccinia virus (VV) – which was long used as vaccine against Smallpox- is the prototype poxvirus, and is the best characterized member of the family (1).

VV entry into cells is a complex process that has been subject to numerous studies. The VV size -approximately 360 nm of diameter- has led to the assumption that its entry into the cell cannot happen through caveolin or clathrin vesicles (3). VV attachment to the cell is thought to occur with glycosaminoglycans and laminin (4, 5). Also, several membrane proteins have been proposed to serve as VV entry receptors, co-receptors or functional signaling ligands (6-8), in a process that leads to VV internalization through fluid phase endocytosis (9-13) followed by membrane fusion. As an alternative route, direct membrane fusion at plasma membrane has also been found to be a way of VV entry to the cytoplasmic space (14-17). Surprisingly, VV entry process varies significantly between different cell lines, virus strains and viral forms (3, 16, 18-20). Taken together, these observations point to the existence of several pathways leading to VV entry, which probably contribute to the broad tropism range of VV.

Genome-wide genetic screens have become an important tool to identify factors that are involved in biological processes. Several technologies, like RNA interference or insertional inactivation have been used to carry out high throughput screens. Recently, tools for the use of CRISPR/Cas gRNA libraries have allowed for a more targeted way of screening genes by inactivation, down- regulation or induction. Several high-throughput screenings using CRISPR/Cas9 technology have been performed to target cellular factors involved in infection viruses like SARS-CoV-2, West Nile virus, Influenza A virus, Human cytomegalovirus and Zika virus among others, revealing many host cell factors involved in viral infection (21-26). In order to unravel the complexity of virus-cell interactions in Poxvirus infection, both RNAi (27-30) and retroviral-mediated insertional mutagenesis DNA (31, 32) have been performed. Overall, the most important hits from those studies include entry factors and cellular genes somehow involved in viral morphogenesis.

We have developed a genome-wide CRISPR/Cas9-based screening using a human genome wide sgRNA GeCKO library that depends on Cas9 expression and delivery of a sgRNA library into cells for target gene deletion (33, 34). The GeCKO library, designed to target all the genes of the human genome was used to obtain a pooled population of single-gene knock-outs that was then subject to Vaccinia virus infection. Through this approach, we have identified B2M as a pro-viral factor related to Vaccinia virus entry. Notably, B2M encodes the well characterized β_2_-microglobulin, a molecular chaperone known to form complexes with multiple partners, including HLA, HFE, FcRn, MR1 (Human Leukocyte Antigen, Hereditary hemochromatosis protein, Neonatal Fc Receptor and Major histocompatibility complex class I-related gene protein). For those proteins, interaction with B2M is needed to reach the cell membrane (35-38). We show herein that inactivation of B2M gene greatly impairs VV infection and viral production by affecting viral entry.

## Results

### Genetic screen reveals a set of hits potentially involved in VV infection

A loss of function, genome-wide screen was carried out by infecting a pooled population of cells harboring single gene KOs. The cell population was obtained by using a lentiviral library of 123,411 sgRNAs covering 19,050 genes and 1,864 miRNAs in the human genome (33). HeLa cells constitutively expressing Cas9 were transduced with the lentiviral-sgRNA library, and passaged for 7-14 days in selective media. Of note, inactivation of genes essential for cell survival or growth are eliminated by this procedure, and therefore the screen is directed to the part of the genome that is non-essential in cell culture, which constitutes about 90% of the cellular genes (39, 40).

We initially screened for gene KOs that could render the cells resistant to VV infection (Fig 1). However, several experiments performed by infecting with VV strain WR under various multiplicities of infection resulted in complete cell death at 48-72 h.p.i. Even low m.o.i. infection resulted in no surviving cells, indicating that no single gene KO was sufficient to completely block infection. This result was consistent with a possible lack of complete resistance to infection by gene inactivation, in agreement with previous results (32). Given that the replicative strain WR was used, we reasoned that even if some degree of resistance would be present in certain cells in the pool, the progeny virus in the cultures could result in the death of initially surviving cells because of a second round of infection. In any event, based on these results, we concluded that probably only partial resistance could be achieved by single gene inactivations.

**Fig 1.**
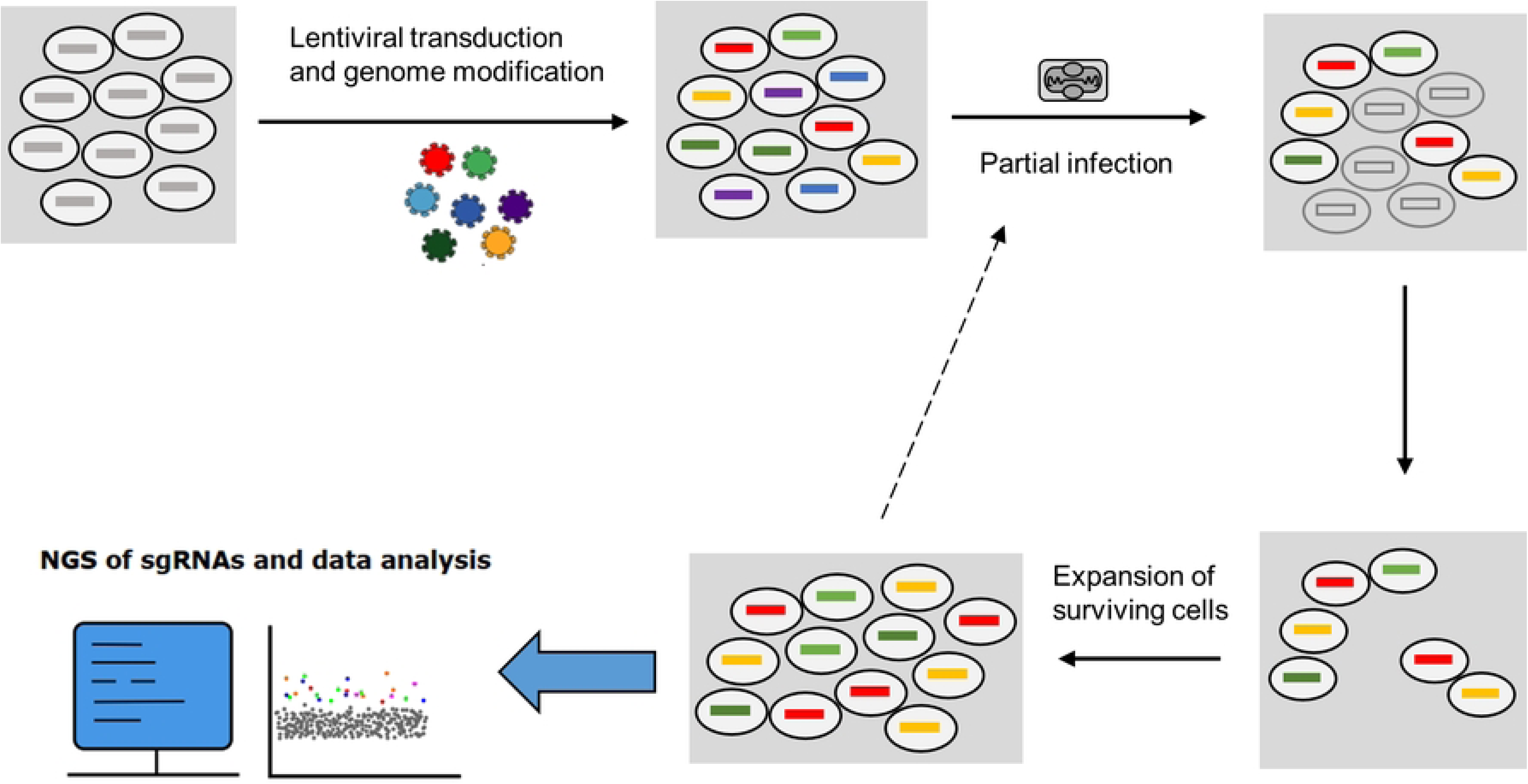
Design of the pooled genetic screen for the identification of VV pro- viral factors. A pool of cells with single gene inactivations was obtained by transducing the GeCKO sgRNA lentiviral library. After infection, surviving cells are analyzed by high-throughput sequencing of the sgRNA region in the integrated lentiviral construct, which is amplified by PCR. Finally, results are analyzed using MaGeCK and ScreenBEAM algorithms for hit determination.

To search for pro-viral genes whose inactivation might lead to partial resistance to infection, we adapted our screen by introducing three main modifications: 1) we aimed at using large cell populations and controlled m.o.i. to infect only a portion of the cells, 2) we performed several consecutive rounds of infection to enhance the enrichment effect and 3) we used viruses with limited replication or production of extracellular virus to prevent death of the surviving cells by infection with progeny virus.

For the ensuing experiments, viruses with limited spread ability in the culture were used. A virus deficient in genes A27L and F13L (V- ΔA27-ΔF13), which has a defect in the production of extracellular virus particles limiting its transmission (41, 42); and a virus deficient in the D4R gene (V- ΔD4), which is blocked in DNA replication of the viral genome and therefore is unable to generate infectious progeny virus (43). In the case of the latter, a complementing HeLa cell line constitutively expressing VV D4 protein (HeLa-D4) was engineered to allow virus expansion in a similar way to previously reported (44). Using those two virus mutants and different m.o.i.s, a number of experiments were completed (Table S1), in which surviving cells were expanded, and analyzed. In parallel, uninfected cells were passaged for two weeks and used as control.

High throughput sequencing results for the sgRNA region amplified by PCR were subjected to bioinformatics analysis. Our computational analysis using MaGeCK (45, 46) algorithm allowed us to identify genes that were enriched. Fig 2A shows the analysis of a particular experiment and Table S2 the results from all experiments. In parallel, passages of non-infected cell cultures pointes to gene inactivations leading to self-enrichment, with similar results to current databases (i.e. DepMap Portal) (47). The degree of enrichment in infected versus non-infected cells led to a refined list of possible pro-viral genes. We obtained a curated hit list selecting those candidates with a False Discovery Rate (FDR) < 0.05 in MaGeCK analysis, which is a narrow confidence threshold. We also used ScreenBEAM algorithm (48) obtaining similar results as with MaGeCK (Table S3). The results from different infection experiments and from the control uninfected cultures is summarized in Fig 2B.

**Fig 2.**
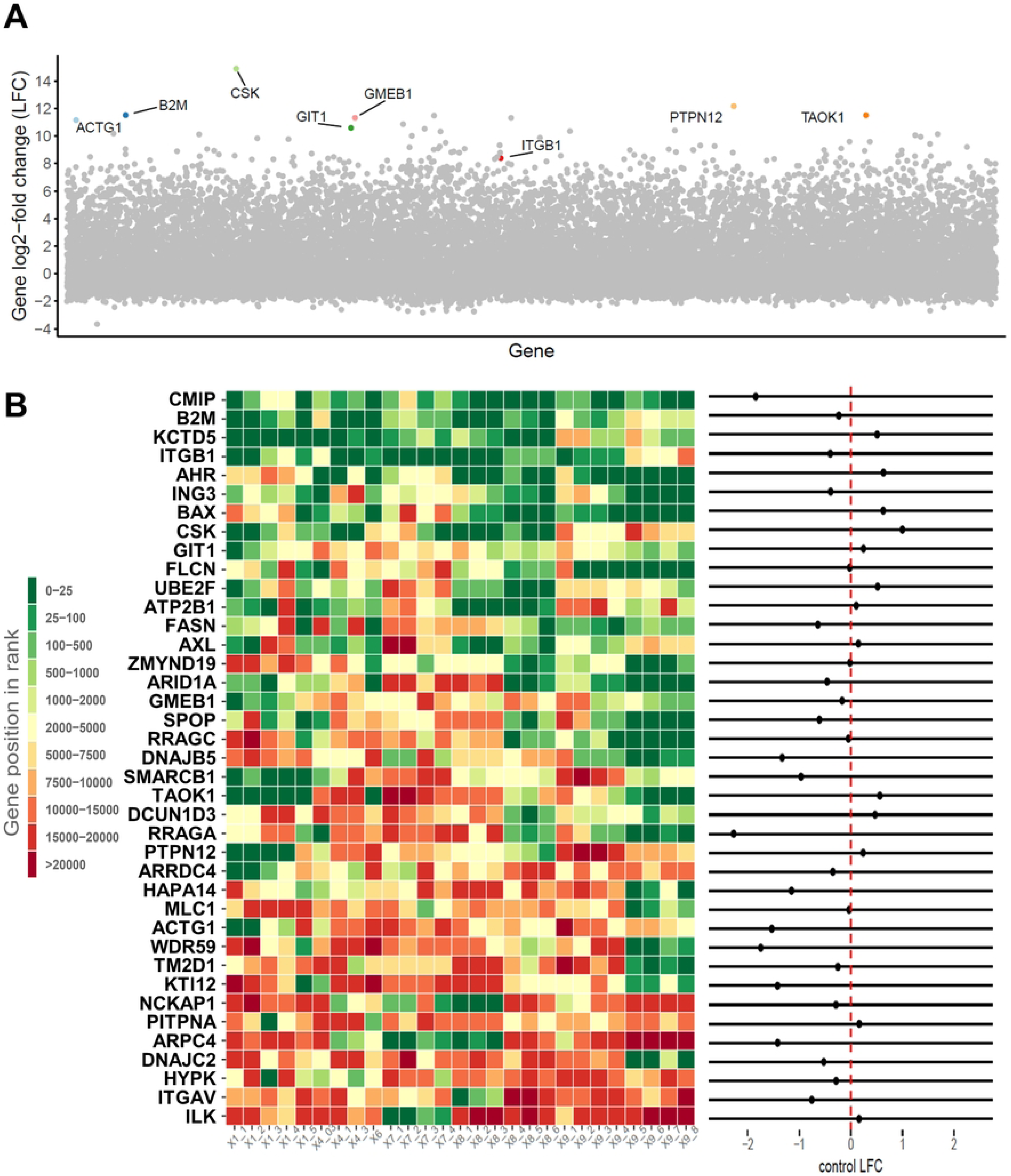
Hit identification. (A) Example of results obtained in one screen experiment analyzed with MaGeCK. Hits with FDR < 0.05 are labelled with gene name tags. (B) Heatmap summarizing the 27 experiments/experiment variations. Genes with FDR < 0.05 in at least 1 experiment are included. The results of enrichment after selection is shown in the left panel as a heatmap representing the gene rank obtained with MaGeCK. For comparison, the right panel shows the log2-fold change for each gene in non-infected control experiment. Dots to the right of the red dotted line indicate enrichment in non-infected cultures. For instance, CMIP KOs are depleted from the pool during passages in the absence of infection, whereas CSK KOs are enriched

### Functional protein networks

Gene hits from multiple experiments were analyzed using the ReactomePA software to identify regulatory pathways (Fig. 3, Fig. 4 and Table S4). Several high significance hits (FDR < 0.05 and LFC > 0) included genes previously described to play a role in the initial steps of VV infection such as *ITGB1* or *AXL* (6, 8), as well as genes encoding proteins involved in their downstream pathways. For instance, integrin alpha subunits and actin cytoskeleton remodeling proteins (*ACTG1*, *WASF2, ARPC2, ARPC3, ARPC4*…) are related to the ITGB1/AXL and AKT activation pathway (Fig 3). Other of our strongest hits is *CMIP*, a poorly studied gene that is downstream-related to ITGB1 pathway (49, 50). Although weaker hits, two annexins were also identified with ScreenBEAM algorithm (*ANXA2* and *ANXA8L2*), as was a member of the S100 family (S100A5) known to interact with annexins (51, 52) (Table S4). Annexins are commonly used to block phosphatidylserine due to its specific binding, and have been shown to reduce VV infectivity when pretreating VV mature virion with Annexin 5 (12).

**Fig 3.**
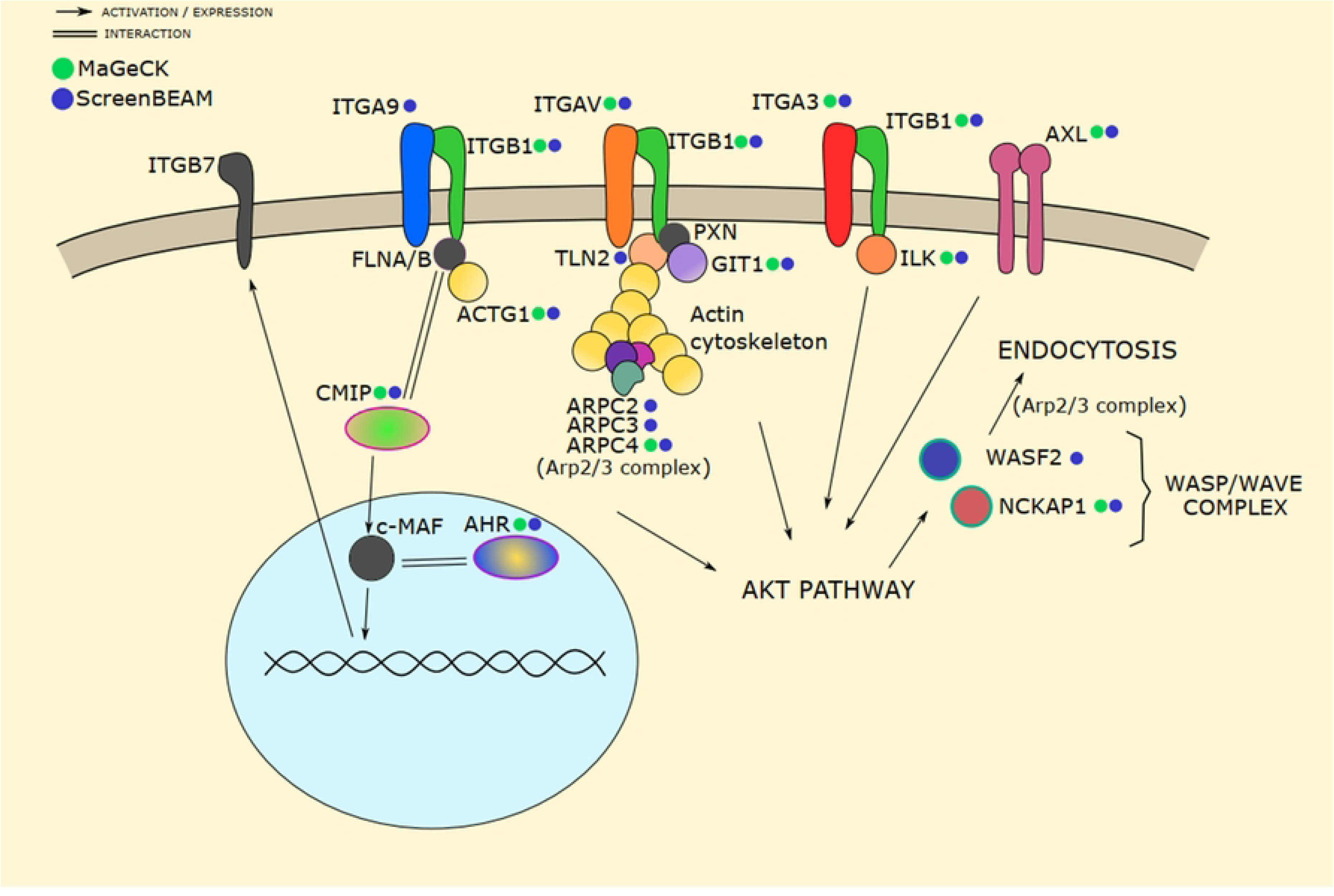
Screen hits related to the ITGB1/AXL activated AKT pathway. Proteins marked with a dot were hits in the screen analysis (MaGeCK: green dots; ScreenBEAM: blue dots). Activation of AKT by ITGB1 signaling pathway results in actin cytoskeleton reorganization, leading to VV endocytosis. CMIP and AHR are new suggested players in the pathway (see main text).

**Fig 4.**
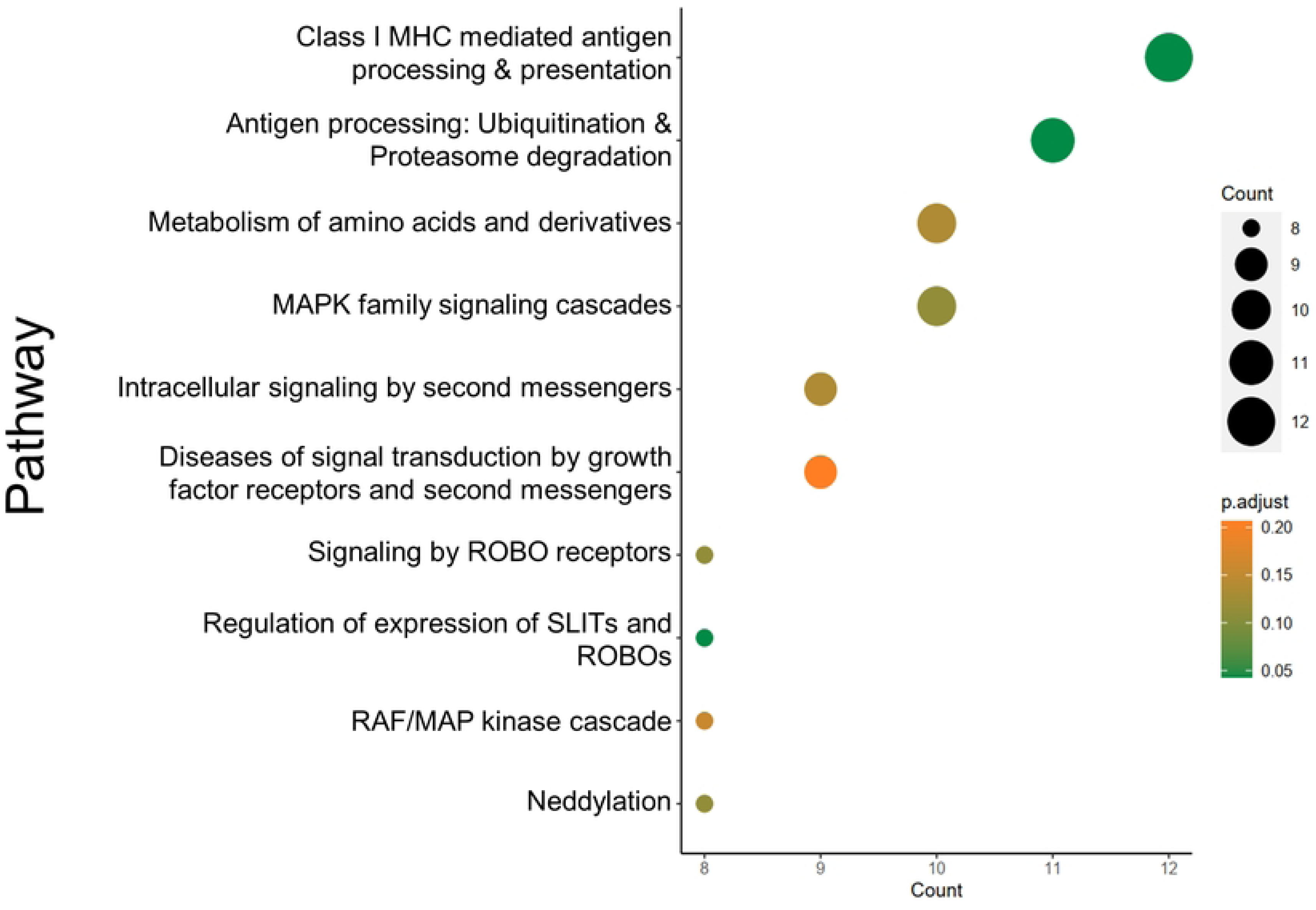
Pathway analysis. Reactome Pathway Analysis was performed using the top 1 % list of genes according to MaGeCK rank. Class I MHC mediated antigen processing & presentation pathway was the most enriched pathway with 12 genes and p-adjust value < 0.05. Genes associated to this pathway by ReactomePA were B2M, UBE4A, CUL3, KLHL22, PSME1, UBE2F, RNF41, PSMC3, TRIM4, PSMA8, TRIP12, ASB17.

Other hits were related to additional pathways known to be important during infection, like the proteasome-ubiquitin pathway. KCTD5 emerged as one of the strongest hits, although it showed some enrichment in uninfected controls. Remarkably, this KCTD5 protein has been proposed to be an adaptor for cullin3 ubiquitin ligases and is known to modulate the activity of Rac1 protein, a GTPase involved in VV internalization (53, 54).

Interestingly, B2M (the gene encoding β_2_-microglobulin) was among the best hits. When analyzing the top 1% genes in the MaGeCK rank by Reactome Pathway Analysis (55) the B2M-related ‘Class I MHC mediated antigen processing & presentation’ pathway was the most prominent (Fig 4 and Table S4). Enriched genes associated to this pathway were B2M, UBE4A, CUL3, KLHL22, PSME1, UBE2F, RNF41, PSMC3, TRIM4, PSMA8, TRIP12, ASB17. All of them except for B2M belong to proteasome/ubiquitin ligases pathway, whereas B2M is more related to post antigen-processing and class I MHC loading stage.

Given that B2M likely represented a novel pathway in VV entry, we reviewed the screening results to check if genes related to B2M were also positive in the screen. Notably, although with lower significance, *TAPBP* was detected in ScreenBEAM analysis (Table S3). Other B2M related genes coding proteins like Antigen peptide transporter 2 (TAP2), Protein disulfide-isomerase A6 (*PDIA6*) and BiP chaperone (*HSPA5*) had good scores, even though their respective FDRs (MaGeCK) or ranking criteria (ScreenBEAM) were below our initial cutoff. Those results reinforce the notion that B2M and its related pathway are bona-fide pro-viral factors as revealed by the survival screen. B2M protein is known to bind a number of proteins that include HLA molecules, HFE or FcRn. We individually tested the scores for those genes but did not find any of those significantly enriched in the screening. Notably, published results on experimental infections of B2M deficient mice reported unexpected levels of resistance to VV (56, 57), phenotypes that now might be reasoned to be influenced by B2M absence.

### *B2M* KO strongly impairs VV infection

The screen results point at B2M as a relevant novel factor in VV infection. To further study its role in VV infection, we obtained HeLa and HAP1 KO cell lines by CRISPR/Cas9-mediated gene-inactivation. Inactivation of B2M gene was confirmed by sequencing of the mutated region in HeLa and HAP1 cells lines, and loss of expression was followed by Western blot and immunofluorescence analysis in HeLa clones (Fig 5A and 5B). Low multiplicity infections (m.o.i. = 0.05) of those clones led to a decrease in progeny virus, indicating an effect of B2M absence on VV infection (Fig 5C). Those results were confirmed in two independent cell clones (2.18- and 1.64-fold reduction). Those results were also confirmed in the background of a second cell line, HAP1.

**Fig 5.**
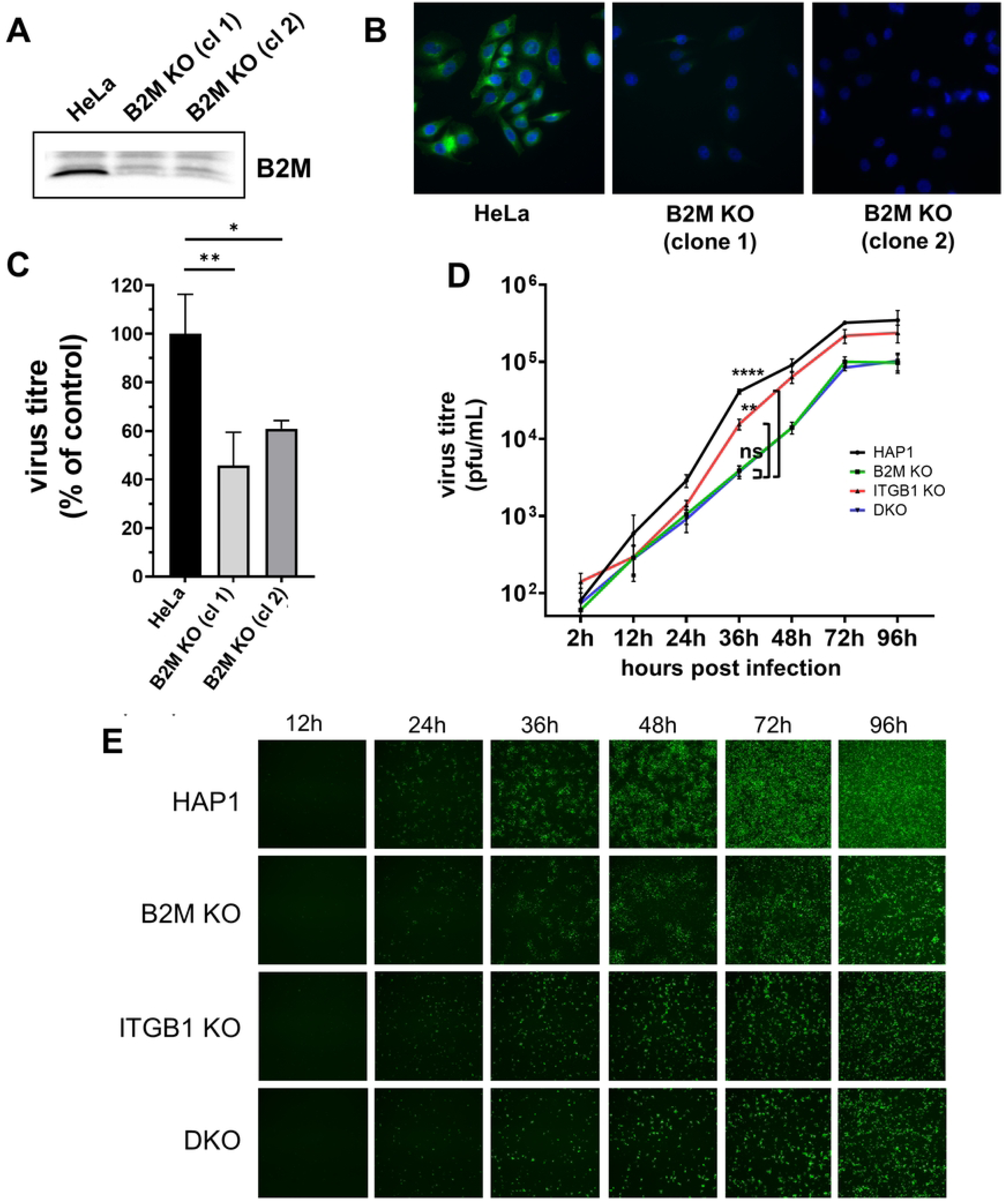
Effect of B2M KO on VV replication. (A) Analysis of cell lysates using anti-B2M antibody. (B) Immunofluorescence images of HeLa and HeLa-B2M KO cells labelled with anti-B2M antibody. (C) Virus production in HeLa and HeLa-B2M KO. Infected cell monolayers (m.o.i. = 0.05) were collected 30 h.p.i. and VV titers were obtained by a standard plaque infectivity assay. Error bars indicate SD of three independent experiments. D and E: HAP1 and derived cell lines were infected at m.o.i. = 0.05. At different time points, a fraction of supernatant was collected to determine extracellular virus production (D). Error bars indicate SD of three independent experiments. In the same cultures, fluorescence images were obtained at different times (E). p-values: **** < 0.0001, *** < 0.001, ** < 0.01, * < 0.05, ns > 0.05.

Since ITGB1 has been described to be involved in VV entry (6), we were interested in studying whether the function of B2M and ITGB1 are overlapping. Infection of ITGB1 KO, B2M KO, and B2M/ITGB1 double KO (DKO) cell clones in the HAP1 cells at low m.o.i. with a GFP-expressing VV revealed a significant reduction associated with B2M inactivation, and a smaller effect associated to ITGB1 inactivation, as measured by both viral production and GFP expression (Fig 5D and 5E). Concurrent inactivation of B2M and ITGB1 genes did not result in enhanced reduction when compared with B2M inactivation alone. We additionally tested if plaque size or number were affected by inactivation of B2M. For this, a GFP-expressing VV was used in a standard plaque assay in the HeLa B2M KO cells. At 24 hours after infection, there was a reduction of 60% approximately in the number of plaques (Fig 6A). In addition, plaque size was clearly diminished (Fig 6B). Overall, these results indicate that VV replication was being negatively affected by B2M inactivation.

**Fig 6.**
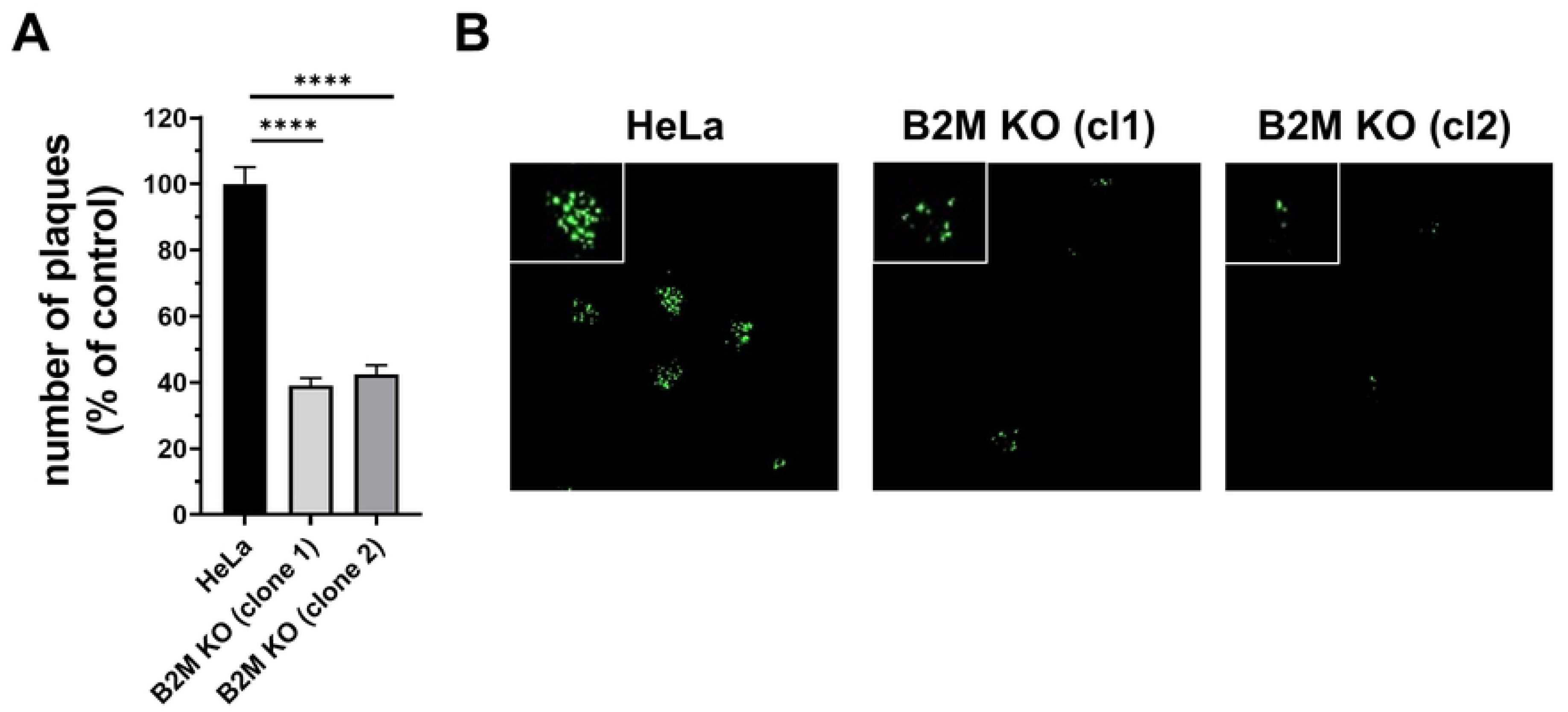
Vaccinia virus plaque assay in HeLa and B2M KO cells. MW6 well plaques with HeLa and B2M KO clones 1 and 2 were infected with 200 pfu/well of VV-GFP. Plaques were counted at 24 h.p.i (A). Error bars indicate SD of three independent experiments. p-values: **** < 0.0001, *** < 0.001, ** < 0.01, * < 0.05, ns > 0.05. (B) Images of representative plaques under the microscope by GFP fluorescence.

### Effect of B2M deficiency on virus entry

We next investigated the stage in the replication cycle which was being affected in B2M KO cells. We anticipated an effect on an early step in the infection cycle, since viruses restricted to the early phase of the cycle (such as the D4-defficient virus) result in complete cell death (not shown). We therefore hypothesized that a block before that stage would be required for cell survival in the screen. Consequently, we focused on the virus entry process. We used an assay based on early luciferase expression (58), where an evaluation of the viral entry can be assessed by the luciferase activity, which is expressed under an early promoter and therefore, immediately after entry but before replication. Virus entry was significantly reduced as a result of B2M inactivation, both in HeLa and HAP1 cells (Fig 7A), indicating that inhibition of infection occurs at an early stage in the virus cycle, before the start of DNA replication.

**Fig 7.**
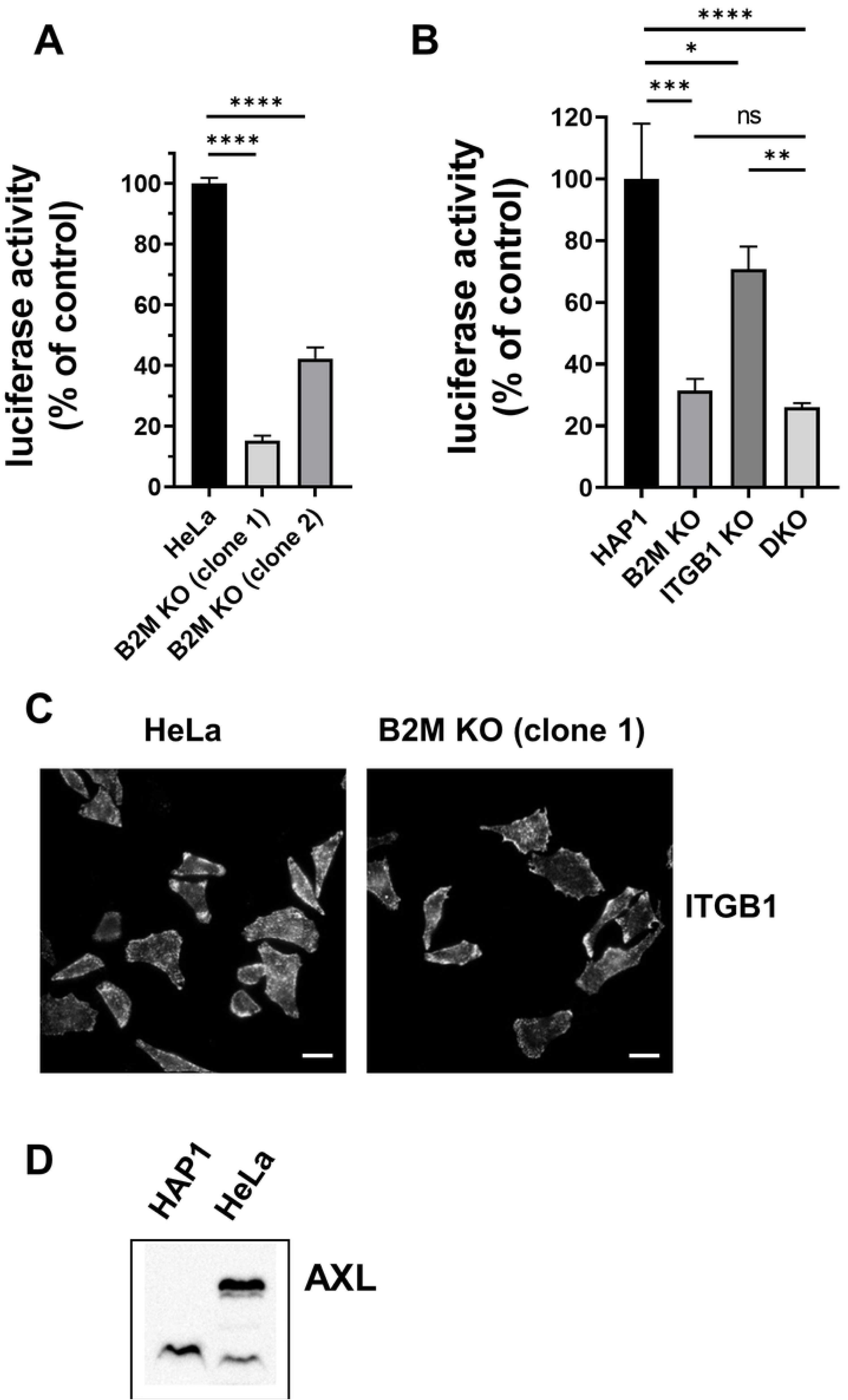
B2M is necessary for an early stage in VV infection. (A and B) Early gene expression was taken as an indicative of virus entry. HeLa (A) and HAP1 (B) cells were infected with VV-Luc at m.o.i. = 0.8 and luciferase activity was measured 3 h.p.i. Error bars indicate SD of three independent experiments. (C) Immunofluorescence of non-permeabilized HeLa and B2M KO cells marked with anti-ITGB1 showing plasma membrane expression and distribution of ITGB1. (D) Absence of AXL in HAP1 cells. Cell lysates of HeLa and HAP1 cells were analyzed by Western blot with anti-AXL antibody. p-values: **** < 0.0001, *** < 0.001, ** < 0.01, * < 0.05, ns > 0.05. Scale bars: 20 µm

Given the reported role of ITGB1 in VV infection, it could be reasoned that B2M inactivation might result in a decrease in ITGB1 levels, which in turn would inhibit VV infection. However, an indication that this might not be the case was that inactivation of B2M was more inhibitory than that of ITGB1 in our experimental setup (Fig 7B). Also, since B2M is required for the transport of several membrane proteins from the ER to the plasma membrane, it could be argued that ITGB1 transport could be affected in the absence of B2M. To assess this possibility, we performed immunofluorescence with anti-ITGB1 antibodies in HeLa B2M KO cells (Fig 7C) Notably, ITGB1 levels in B2M KO cells were similar to that in parental HeLa cells, indicating that ITGB1 was transported efficiently to the plasma membrane independently of B2M. Therefore, we concluded that B2M KO effect on VV infection is not mediated by a decrease of ITGB1 in the cell surface.

An additional screening hit, AXL, has also been implicated in VV internalization (8, 12, 32, 59). However, we do not consider likely that the B2M effect on virus entry is mediated by AXL absence, since HAP1 cells do not express AXL (60). This result was confirmed by Western blot, where the expression of AXL is detected in HeLa but not HAP1 (Fig 7D). This result is in agreement with AXL being an enhancer, albeit not an essential factor for infection (8, 61).

### *B2M* inactivation does not affect VV binding but delays internalization

In order to test if virus attachment was affected in *B2M* KO cells we performed a binding assay in HeLa *B2M* KO cells. To measure binding, purified VV particles incorporating a fluorescent protein fused to a virion protein constituent (VV A4- cherry) were incubated with cells and maintained at 4 °C to avoid virus internalization, and virus particles bound to cells were quantitated by confocal microscopy (Fig 8A and 8B). Results showed no significant differences between control and *B2M* KO cells indicating no correlation between B2M presence and VV attachment.

**Fig 8.**
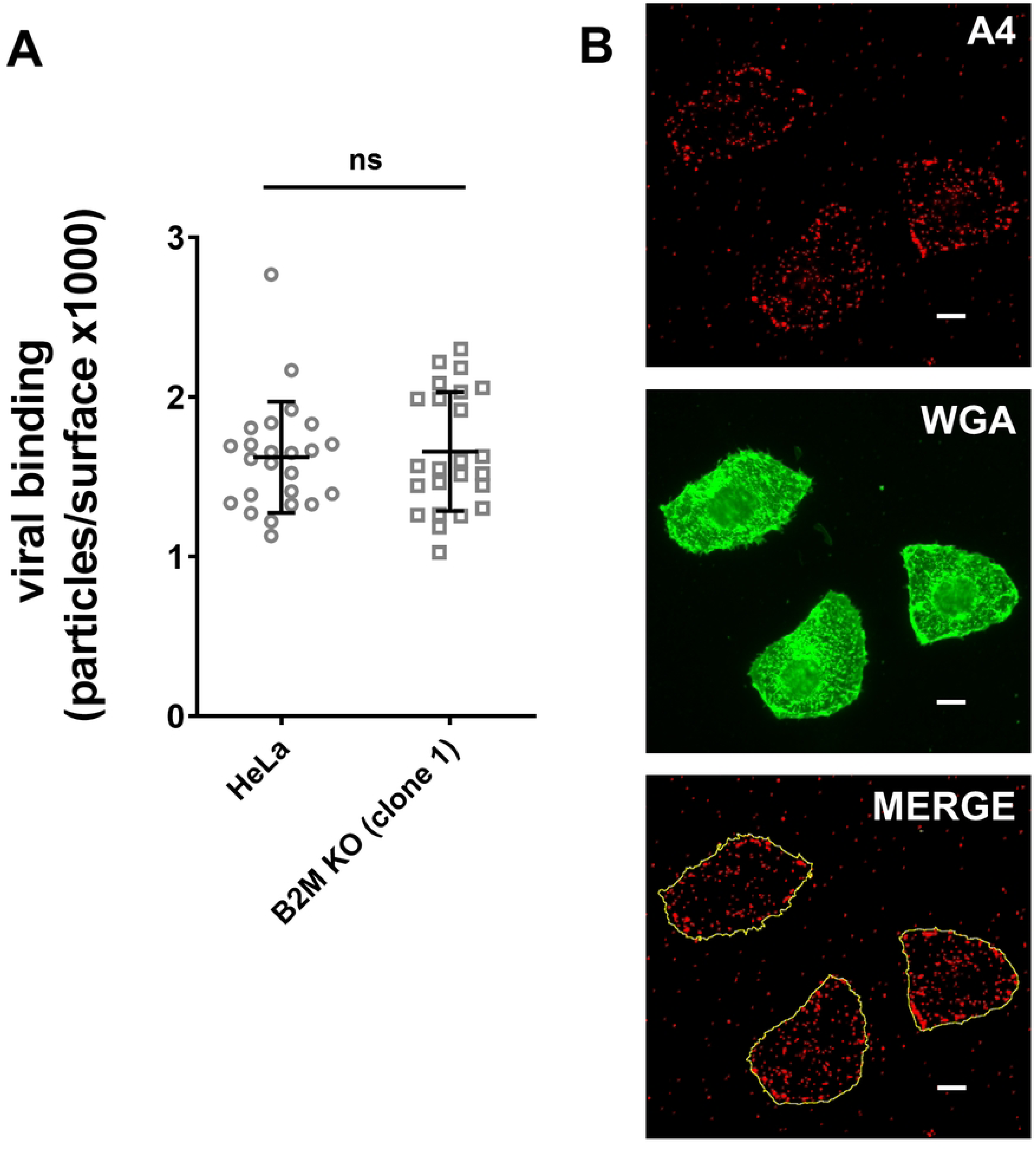
VV binding is not affected by B2M absence. (A) HeLa and B2M KO cells were infected with VV A4-cherry at m.o.i = 30. After 1 h of adsorption at 4 °C, cells were washed with PBS and viral particles bound to the cell were counted. To standardize measurements data is given in relation to cell surface area. (B) Example showing the labeling with WGA-488 to visualize the cell surface. p-values: **** < 0.0001, *** < 0.001, ** < 0.01, * < 0.05, ns > 0.05. Scale bars: 10 µm.

We next analyzed the effect of B2M absence on the efficiency of VV internalization. For this, after the initial virus adsorption period, cells were incubated at 37 °C to allow virus internalization and therefore, internalized virus particles were quantitated at different times (Fig 9A). We used VV A4-cherry (for total virus) and an antibody targeting the VV Intracellular Mature Virion surface protein A27 to detect those viral particles that were not internalized (Fig 9B). Virus internalization occurred progressively over time in HeLa cells, similarly to previous reports (9). Comparison with HeLa cells deficient in B2M showed a consistent decrease in the internalization rate in *B2M* KO cells, an effect that was more pronounced at the intermediate tested times (Fig 9A). These results suggest that a delay in virus internalization exists as a result of B2M inactivation, although some virus particles are eventually able to reach the cell interior.

**Fig 9.**
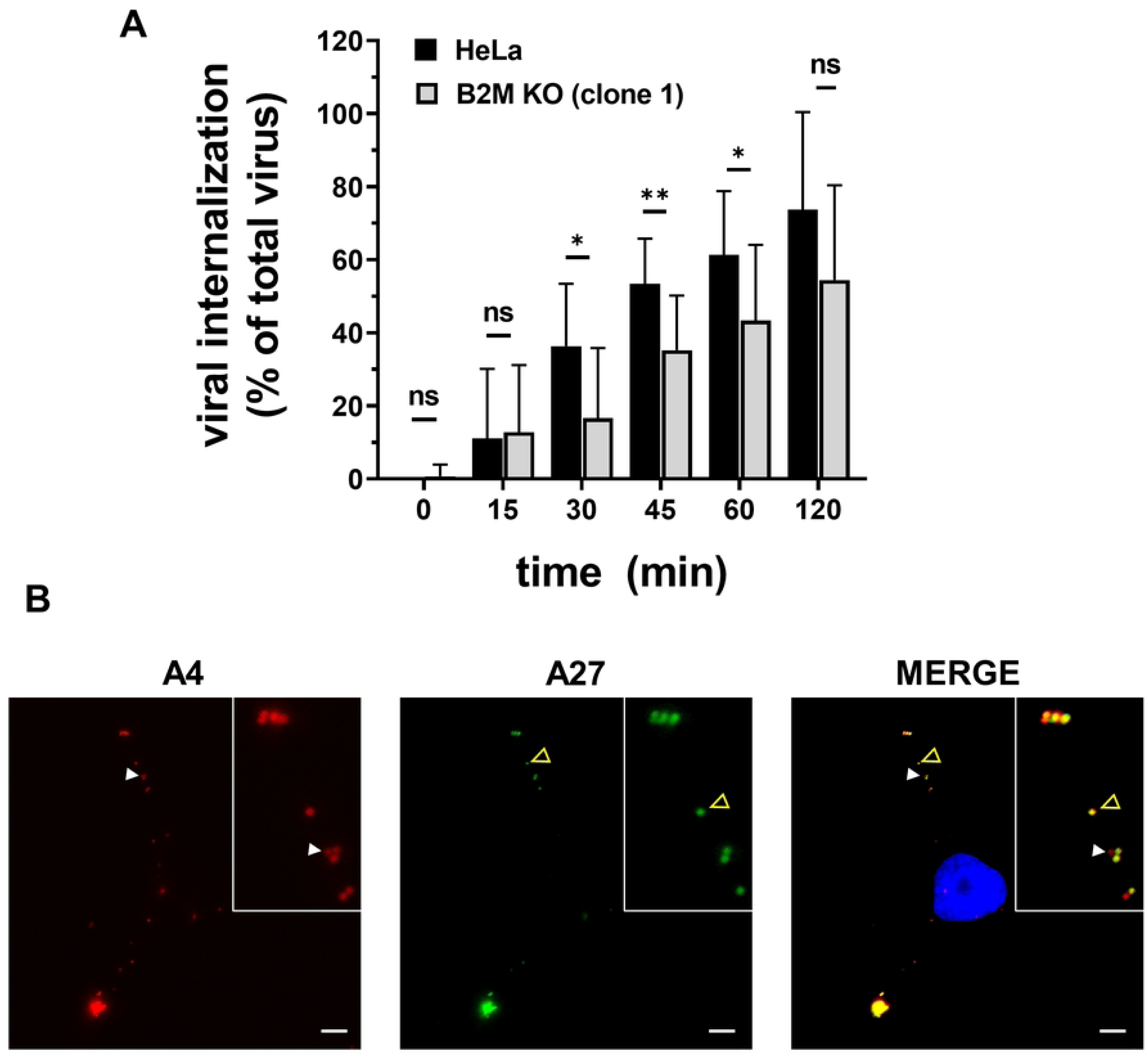
B2M gene inactivation affects VV internalization. (A and B) HeLa and B2M KO cells were infected with VV A4-cherry at m.o.i. = 4 during 1 h at 4 °C to avoid internalization. Cells were incubated for different times at 37 °C and non-permeabilized cells were stained with antibody to VV-A27 (green) to visualize cell surface virus particles. White arrow-heads mark internalized virions (only red in merged image) and yellow-empty arrow-heads indicate non-internalized virions (green or yellow). Cell nucleus was labeled in blue by Hoechst staining. Internalized virions were quantified for each time point showing significant differences between HeLa and B2M KO cell lines. p-values: **** < 0.0001, *** < 0.001, ** < 0.01, * < 0.05, ns > 0.05. Scale bars: 5 µm.

### VV colocalization with B2M

As the specific role of B2M in early VV infection is related with the early steps of infection, we tested the degree of colocalization of B2M and VV at cell membrane. Since viral entry is a sequential process in which different protein-protein interactions may occur over time (62, 63), different time points were included in the experiment. After adsorption of fluorescent VV A4-cherry particles at 4 °C, cells were stained with antibodies specific for B2M and ITGB1. Confocal images of non-permeabilized cells revealed a partial colocalization of B2M and attached virus particles (Fig 10A). We found approximately 50 % of VV particles colocalizing with B2M at 0, 15, 30, 45 and 60 min post infection (S2 Fig). To determine if these colocalization results were consistent, we estimated VV/ITGB1 colocalization, that has been described to occur, at least in HeLa cells at 0 min (6). Our results showed that 38 % of adsorbed VV particles colocalized with structures containing ITGB1. Also measured that 24% of B2M- positive structures were also labeled with ITGB1, and that 20% of ITGB1- positive structures were also labeled with B2M. Those results indicate only partial colocalization of the two proteins with each other, and with the infecting VV particle, and show a substantially differential pattern for B2M and ITGB1 (Fig 10B).

**Fig 10.**
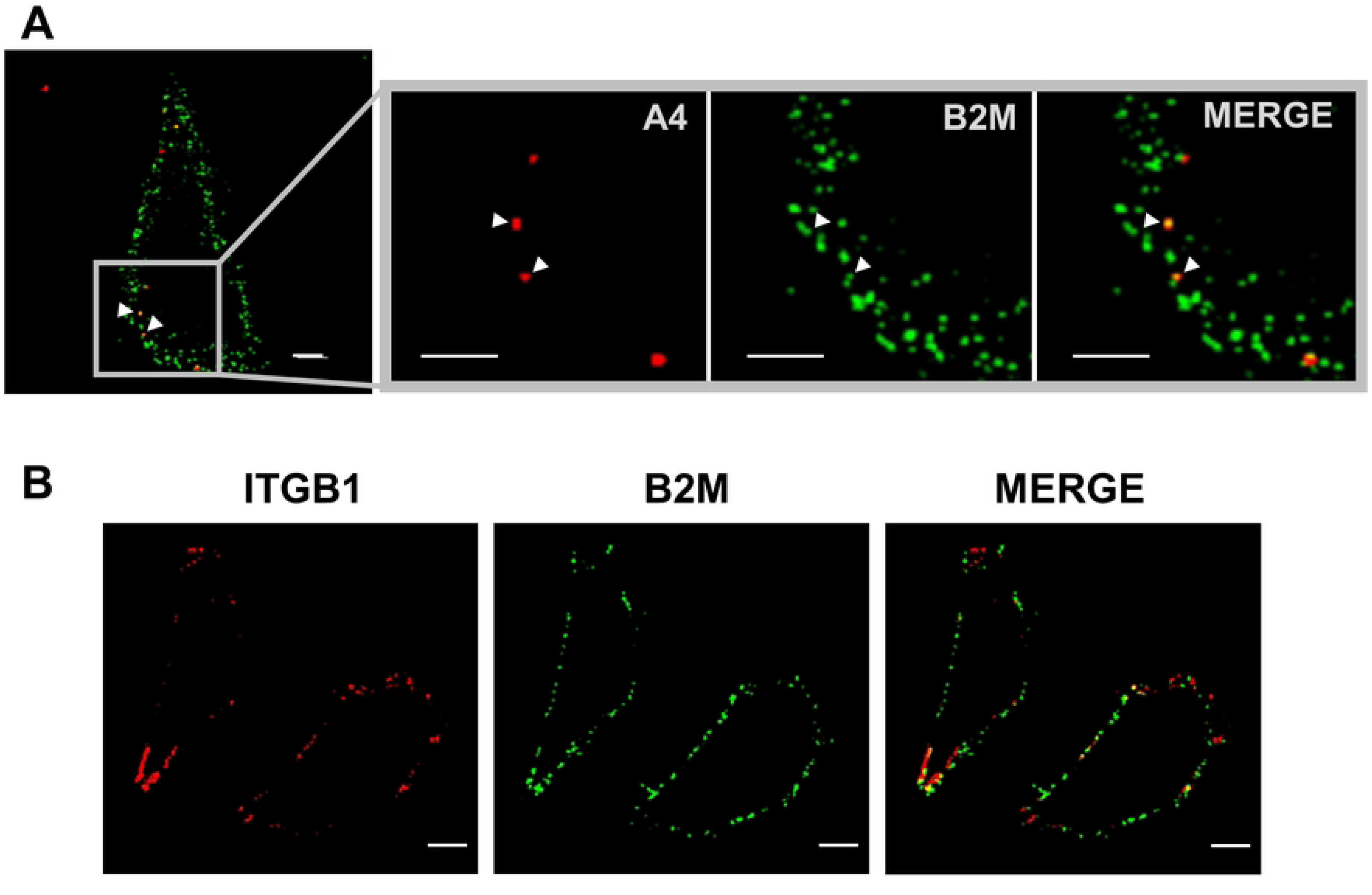
VV, B2M and ITGB1 partial colocalization. (A) HeLa cells were infected with VV A4-cherry (red) for 1 h at 4 °C and unbound virus was removed by washing (m.o.i. = 5). Cells were stained with anti-B2M antibody (green) to analyze colocalization. Colocalization of VV with B2M is indicated with white arrow-heads. (B) Non permeabilized HeLa cells were stained with anti-B2M (green) and anti-ITGB1 (red) to analyze colocalization. Scale bars: 5 µm.

Finally, we wished to determine if B2M was able to block virus infection by interacting directly with the virus particle. For that, purified VV particles were incubated with soluble B2M and then used for infection. In those experiments, no significant differences were found (S3 Fig). On the other hand, a blocking assay was also performed pretreating the cells with different concentrations of anti-B2M antibody, followed by VV infection but we did not observe any significant blocking activity (S4 Fig).

## Discussion

### CRISPR/Cas9 genetic screen

The interactions between VV and the host cells have been a subject of considerable research. In this work, we have used CRISPR/Cas9 technology to perform a pooled, genome-scale loss-of-function screen with the aim of identifying cellular genes necessary for VV infection. From the design of our screening, it should be noted that we are only testing the non-essential gene set, since after subjecting pooled KO cell population to six passages, the cells in which essential functions are inactivated are depleted from the pool. Likewise, an enrichment is observed for some KOs in genes that perform functions related to the control of cell proliferation and apoptosis. This observation is common in screens of cellular genes (47).

The experimental approach to this problem was complicated by the considerable versatility and adaptability of Vaccinia infection. In fact, in our initial experiments, all cells showed cytopathic effect and suffered cell death over time, regardless of the multiplicity of infection used in the experiment. Failure to obtain even a single surviving cell suggests that a single KO mutation is not enough to confer complete resistance in the cells. This could be due to a number of different reasons. A first explanation might be that the virus is able to enter the cell using different, non-overlapping molecular pathways. Second, even if there is partial resistance, it may be difficult to isolate a few resistant cells within a large cell population of virus-producing cells. Indeed, since the bulk of the cells would remain susceptible they would generate high loads of progeny virus that may overcharge and kill the partially resistant cells. In this respect, it has been shown that even inactivated virus can induce cell death (64). In similar approaches, high multiplicities of infection resulted in the death of the whole cell population, a difficulty already reported by Luteijn et al. (32).

In view of these considerations, we modified the screening protocols to exert a gradual enrichment of partially resistant cells. Those modifications included infection with viruses that have decreased spread or limited replication, and adjusting the multiplicity of infection to modulate the selective pressure. The screening design was directed at maintaining a controlled selective pressure in the form of virus-induced cell death in the cultures during extended periods. Importantly, this may favor the enrichment of cells harboring gene-KOs leading to enhanced cell growth, as we have seen over extensive passage of uninfected samples. Therefore, enrichment in our experiments may stem from increased resistance to infection or from faster cell growth. This was taken into account when analyzing the possible hits in the screening by comparing sgRNA changes in infected and control uninfected cultures (Fig 2B).

We have observed that viruses blocked in DNA replication, such as the V-ΔD4 virus mutant, efficiently promote cell death even though replication cycle is not completed. This indicates that cell survival is not possible after the early stage of virus infection is reached. Because our screening was based on cell survival, it not surprise that it was preferentially directed to the cell entry process. Vaccinia virus and Poxviruses in general are known to have a complex entry process into host cells, where they first bind in an unspecific manner to extracellular matrix components such as glycosaminoglycans (GAGs) and laminin, followed by a more specific attachment step (4, 6, 63). During this process, signal activation pathways are triggered in the cell, leading to the internalization of the virus through fluid phase uptake and -to a lesser extent- direct fusion at plasma membrane (7, 12, 58). With our screen approach, we identified several potential gene candidates that may play a role in early stages of infection. Statistically significant hits like *ITGB1* and *AXL*, as well as some of their downstream pathway proteins such as Akt-pathway-related proteins, the small GTPases –RhoA, Rac1, Cdc42-, or actin cytoskeleton remodeling proteins (6, 8, 65-67) were obtained. Additional gene hits were also related to these pathways. For instance, c-Maf activating protein (CMIP) was one of the strongest hits and is thought to participate in several cellular processes, including integrin expression through c-Maf activation, actin cytoskeleton remodeling and the Akt route (49, 50, 68). Another hit was AhR (*AHR*), which is known to be bound to c-Maf (69, 70). We suggest that CMIP may have emerged as a hit in our screen due to the concurrence of the different actions where this protein is involved (Akt pathway, cytoskeleton, integrin expression…) (50, 71, 72). Because CMIP is a poorly studied gene, it will be of interest to study its potential function in VV infection. Overall, this study contributes to expand the big picture of the likely cell - VV interactions during the first steps of infection.

### β_2_-microglobulin gene (B2M)

Unexpectedly, B2M gene appeared as one of the strongest hits in the screen, implying B2M protein is an important player in VV infection. B2M is a molecular chaperone that is expressed in most mammalian cells and binds several protein ligands such as MHC-I, HFE or FcRn before their transport to the cell surface. We used different approaches to determine the role of B2M in VV infection. First, we found that B2M deficiency leads to a decrease in viral plaques, and a lower viral progeny titer. Second, B2M role is relevant during early infection stages, since early gene expression was clearly diminished (Fig 7). More precisely, we demonstrated that B2M absence does not affect VV binding but it does decrease internalization (Fig 8 and Fig 9). Although internalization differences were around 25 % between *B2M* KO and normal cells, these differences were much greater in early expression luciferase assays (around 70%). Several explanations may account for the difference: for instance, the existence of an unproductive internalization pathway in *B2M* KO cells, not leading to early transcription. Also, the internalization delay and defect in *B2M* KO cells may be indicative of the inefficacy of VV virions finding their signaling ligands at cell surface, therefore abrogating them to the unproductive pathway above mentioned.

ITGB1, which has also been shown to be involved in VV entry, was also a significant hit in the screen,. However, several results point to B2M as acting independently of ITGB1. First, the effect of the inactivation of B2M in VV infection in HAP1 cells was stronger than that observed ITGB1 KO. Furthermore, the depletion of B2M in the HAP1-ITGB1-KO line also resulted in a clear reduction of VV infectivity. This result shows that in the absence of ITGB1 and AXL (the latter not being expressed in HAP1 cells), B2M is still necessary for VV infection. Those results can be explained as B2M acting as a key player in alternative virus entry pathway.

Pathway Analysis of the data from screening experiments showed that Class I MHC mediated antigen processing & presentation pathway was the most enriched one, including many proteins related to the proteasome and ubiquitin system. In this respect, a screen using Monkeypox virus also pointed to several enriched ubiquitin genes and other B2M-related genes, such as TAPBP and CALR (31, 32). The ubiquitin system has been extensively studied for its importance during infection, and its relationship to B2M is hence intriguing in view of our results.

At this point, there is no clear mechanism to account for B2M role during infection. B2M requirement for VV infection might be the result of a direct involvement of this protein in the entry process. However, given that B2M forms protein complexes with a number of membrane proteins, one interesting possibility is that one of those complexes might act as a VV receptor or internalization factor. Nonetheless, we have analyzed our screening results searching for these B2M related genes, and we found no clear results. Gene redundancies, in particular regarding HLA genes, may obscure the screening results, that have the limitations imposed by the fact that only single gene inactivations are tested. Current experiments are directed at exploring if any of those B2M heterocomplexes is indeed involved in virus entry.

The known function of B2M does not offer an evident link with a mechanism of virus entry. However, its function in the initial steps of VV infection is not a singular case. Intriguingly, some similarities exist with Coxsackievirus A9 (CAV9) -an enterovirus B- whose internalization is also mediated by B2M (73-75) and which also makes use of at least two subunits of integrin; beta 6 and beta 3. Furthermore, CAV9 entry is mediated by dynamin, as is also the case with VV (76). Recently, it has been found that the effect of B2M for Enterovirus B entry can be explained by the need of B2M in mediating the transport to the membrane of its receptor, the Human Neonatal Fc Receptor (FcRn) (77, 78). Expanding the similarities to the case of CAV9, we obtained ASAP1 as a hit (Table S3), a GTPase activating protein involved in Arf1, Arf5 and Arf6 activation and cytoskeleton remodeling (79), the latter of which being known to mediate endocytosis for some viruses, like CAV9 (74). As authors suggested, virions might undergo an unproductive entry pathway in B2M absence (74), a hypothesis that is consistent with our results. Due to the several similarities in cellular requirements of CAV9 and VV entry, it is likely that one of the entry mechanisms for VV is shared by CAV9. In any event, more research is needed to further clarify the details of the process.

One last interesting aspect of B2M involvement in virus entry is the relevance of B2M product (β-2 Microglobulin) in the immune response. The involvement of B2M as a pro-viral factor might have multiple consequences regarding the pathogenesis and immune response in animal infection. Interestingly, two reports in the literature found that B2M-defficient mice were remarkably resistant to VV infection, despite the lack of cell-mediated immunity in those animals as a consequence of MHC-I deficiency (56, 57). This result suggests the B2M deficiency has also a VV blocking effect, even though those mice have a T cell immunodeficiency.

Overall, this report adds a new player to our picture of the complex interplay between cell and virus in the VV entry process. Delineation of the protein interactions and pathway activations taking place in this process will undoubtedly involve multiple players yet to be discovered.

## Materials and Methods

### Cells

HeLa (ATCC CCL-2) cells were grown in Eagle’s Minimal Essential Medium (EMEM) containing 7 % fetal bovine serum (FBS). BSC-1 cells (ATCC CCL-26) were grown in Eagle’s Minimal Essential Medium (EMEM) with 5 % fetal bovine serum (FBS). HAP1 and its derived lines (*B2M* KO, ITGB1 KO and double B2M/ITGB1 KO (DKO)) were purchased from Horizon Discovery and were grown in Iscove’s Modified Dulbecco’s Medium (IMDM) containing 7 % fetal bovine serum (FBS). HEK-293T cells were grown in Dulbecco’s Minimal Essential Medium (DMEM) with 7 % fetal bovine serum (FBS). All media were supplemented with 0.1 mg/mL penicillin, 0.1 mg/mL streptomycin, 2 mM L-glutamine (BioWhittaker). Cells were grown in a 5 % CO_2_ incubator at 37 °C.

### Viruses

Vaccinia virus (VV) Western Reserve (WR) strain was obtained from the American Type Culture Collection (ATCC VR-119) and routinely propagated in BSC-1 cells. Previously described viruses V-GFP (recombinant WR expressing GFP) (80), and V-ΔA27ΔF13 (recombinant having the A27L and F13L genes deleted) are already described (41).

Virus infections were performed in media containing 2 % FBS. Viral titers in cell lysates obtained after disrupting the cells by three rounds of freeze/thawing were determined by the standard VV plaque assay in BSC-1 cells (81).

### Isolation D4R expressing cells

Complementing cell lines 293-D4rR and Hela-D4rR were obtained by inserting the VV D4R gene in the genome of the parental cells using the Flp-in T-REx™ system from Invitrogen. The integrative plasmid pCDNA-FRT-TO-D4R was transfected in the cell line HeLa Flp-In™ T-REx™ (R71407) or 293-FITR- Flp- In™ T-REx™ (R78007) and subsequently cell clones were selected for hygromycin resistance. A bicistronic construct was inserted, placing the VV D4R gene downstream of a tetracyclin-inducible promoter and the TagRFP gene downstream of an IRES sequence. Clones generated from 293 and HeLa were chosen by monitoring the appearance of red fluorescence after tetracycline induction. The efficacy of cell line complementation was tested by plaque phenotyping with the V-ΔD4L virus.

### Generation of recombinant and mutant vaccinia viruses

Recombinant virus V-A4-cherry was generated as previously described (76). Briefly, BSC-1 cells (6-well plate format, 10^6^ cells/well), were infected at a multiplicity of infection (MOI) of 0.05 PFU/cell with virus WR. After 1 h viral adsorption, virus inoculum was removed and cells were then transfected with 2 μg of plasmid pm-cherry-A4L (kindly provided by Wen Chang) using FuGeneHD (Promega) according to the manufacturer’s recommendations. After 2 to 3 days, cells were harvested, and recombinant virus was isolated by three consecutive rounds of plaque purification. Viral stocks were generated in BSC-1 cells and viral titers were determined by standard plaque assay (PFU/mL).

V-ΔD4L virus was isolated following the protocol already described (43, 44) by infection/transfection of BSC-1 cells with WR virus and plasmid pΔD4R-dsRed. pΔD4R-dsRed contains the dsRed gene downstream of a synthetic early/late promoter between two DNA segments acting as recombination flanks for the D4L locus. At 72h the culture was collected and several successive rounds of plaque purifications in BSC-1 cells were performed by selecting large red plaques (representing single recombinants). The resulting virus was amplified in the complementing cell line expressing the D4R gene (HeLa-D4R). The amplified stock was plaque purified in the HeLa-D4R line. The size phenotype for each clone was checked by plaquing in parallel in the BSC-1 line and the HeLa-D4L line. Virus clones that produced large plaques in the HeLa-D4 cell line and single infected cells in BSC-1 were selected.

V-e.luc VV recombinant expressing firefly luciferase under the control of an early promoter was constructed by inserting an early-Luc cassette in plasmid pA-E-Luc into the A27L locus using a plaque selection procedure (41).

### Lentiviruses

Lentivirus stock production was performed by cotransfection of HEK293T cells with the lentiviral plasmid with the packaging plasmid psPAX2 and pseudotyping plasmid pCMV-VSV-G. Plasmid psPAX2 was a gift from Didier Trono (Addgene plasmid # 12260; http://n2t.net/addgene:12260; RRID:Addgene_12260). pCMV-VSV-G was a gift from Bob Weinberg (Addgene plasmid # 8454; http://n2t.net/addgene:8454; RRID:Addgene_8454). For sgRNA expression, plasmids lentiGuide-Puro (Addgene plasmid # 52963; http://n2t.net/addgene:52963; RRID:Addgene_52963) or lentiCRISPR v2, (Addgene plasmid # 52961; http://n2t.net/addgene:52961; RRID:Addgene_52961) provided from Feng Zhang, were used. For stock titer determination, HeLa cells on MW6 plates were infected with successive dilutions of the lentiviral particle stock and incubated with the corresponding selective medium (blasticidin 5 μg /mL or puromycin 1 μg/mL). After cell clones were visible at the microscope, titer was calculated from the number of resistant clones.

### Isolation of Cas9 expressing HeLa cells

LentiCas9-Blast plasmid was a gift from Feng Zhang and was obtained from Addgene (Addgene plasmid # 52962; http://n2t.net/addgene:52962; RRID: Addgene_52962). HeLa cells grown on MW6 wells were transduced with different dilutions of a lentiviral stock obtained from LentiCas9-Blast. After applying selection with blasticidin (5 μg/mL) for 10 days, cell clones were visible at certain dilutions from which lentiviral titers were estimated. Cells transduced with an m.o.i. of 3-5 were cloned by limit dilution in MW96 wells and individual clones were harvested and expanded for further analysis. Cas9 expression was verified by immunofluorescence and western blotting using anti-flag antibody. A cell clone with the highest Cas9 expression was termed HeLa-Cas-ML, expanded and used for subsequent experiments.

### CRISPR Knock-Out (GeCKO) v2.0 libraries: lentivirus stock and derived cell lines

The Human GeCKO v2.0 library, a gift from Feng Zhang, in the two lentiviral vector version (Addgene 1000000049), was used (33, 34). The lentiviral library LentiGuide-puro consists of specific sgRNA sequences for gene knock-out covering the gene set of the human genome. The library is supplied as two half-libraries (A and B). When combined, the libraries contain 6 sgRNAs per gene (3 sgRNAs in each library).

HeLa-Cas-ML cells were transduced with lentiviral libraries A and B separately and selected with puromycin to obtain a cell mixture called HeLa-Mix. For each library, cells in 4 subconfluent T150 flasks were transduced with approximately 2 x 10^7^ lentiviral particles. At 24 h, medium was replaced by fresh medium containing 1 μg/mL puromycin. After three days the cells were harvested, pooled, and subcultured to 8 T225 flasks. Subsequently, cells were transferred to 24 T225 flasks maintaining the selective medium at all times for 10 days and constituting the pass 0.

### Genetic Screen

GeCKO KO cell pools in 2 T225 flasks (about 10^8^ cells) were infected in each condition. Cells were maintained in EMEM-7% FBS except during virus adsorption, which was carried out in EMEM containing 2% FBS. Uninfected control cells were maintained in culture for 8 subcultures before analysis. Detached cells from the Infected cultures were removed regularly. Infection was monitored by detecting infected cells by fluorescence (red fluorescence for V- ΔD4L and blue fluorescence for V-ΔA27ΔF13). To exert selective pressure in the cultures in the form of VV infection, cells were occasionally re-infected with fresh inoculum. If cultures reached confluence, they were subcultured into new T225 flasks. To avoid population bottlenecks in the sgRNA population, cell dilutions at each passage were never greater than 4-fold. The last passage was allowed to reach confluence and the cells were harvested and prepared for sgRNA quantitation in the surviving population. The conditions under which the infection experiments were performed are summarized in S1 Table.

After the selective process, cellular DNA was extracted and the lentiviral region containing the sgRNA sequence was PCR amplified and sequenced. For DNA purification cell samples were suspended in 500 μL of lysis buffer (10 mM Tris (pH 8.0), 10 mM EDTA, 10 % SDS and 100 μg/mL proteinase K) and incubated 3 h at 55 °C or o/n at 50 °C. Samples were then extracted with an equal volume of phenol twice, then 3-5 times with a volume of phenol: chloroform: isoamyl alcohol (25: 24: 1) and finally once with a volume of chloroform: isoamyl alcohol (24: 1). DNA was then precipitated by adding one-tenth volume of 3 M sodium acetate and two volumes of ethanol and, cooled to -20 °C for 1 h followed by 30 min centrifugation at 4 °C and 13000 rpm. The pellet was resuspended in 300 μL TE with RNAase mix and frozen at -20 °C for storage.

Amplicon for sequencing was derived from cellular DNA by three successive nested PCRs. PCR reactions were performed with Biotools thermostable DNApol enzyme. The first PCR was performed on all the purified DNA as template, by setting up multiple reactions containing each 5 μg of DNA in 100 μL of reaction volume, with 5’P1, 3’P1 oligonucleotides and 15 amplification cycles. After the first PCR, all reactions were pooled and a second PCR was performed in four-fold less reactions with a 1:20 dilution of the first as template in 100 μl of reaction with oligonucleotides 5’P2, 3’P2, and 20 amplification cycles. The third PCR was performed similarly (1:20 template, 1:4 number of reactions, 20 cycles) to incorporates the sequences and the barcodes necessary for Illumina sequencing. Each sample carries a different combination of R and F oligonucleotides in order to multiplex the Illumina sequencing run. Sample libraries were equimolarly mixed and the resulting pools quantified by dsDNA fluorescence (Quantifluor, Promega) and qPCR (KAPA Library Quantification kit, Roche). Single-end sequencing (76 bases) was performed in a NextSeq 500 (Illumina) sequencer or in a Hiseq (Illumina).

### Statistical analysis and screen data analysis

Control and infected simples read counts were analyzed with MaGeCK test function (45) and ScreenBEAM algorithm (48). MaGeCK log2-fold change (LFC) scores of each gene were used for data visualization using R coding language. A cutoff of FDR < 0.05 was applied for the heatmap representation to select the best hits of each screen experiment. For ScreenBEAM analysis, the 20 best hits were selected from each experiment following z-score criteria. Genes with less than 3 successful sgRNAs as well as gene knockouts enriched in control were filtered out. ReactomePA (R package) was performed using the top 1 % gene list from MaGeCK output. Statistical analysis were performed using Student’s *t* test and one-way ANOVA (when more than two group comparisons) using *Bonferroni* correction method (GraphPad Prism 8 and RStudio 1.2.5033).

### Isolation of KO cell clones

HeLa B2M KO lines: To obtain a B2M KO cell line, pLentiCRISPRv2 was digested with BsmBI, and a pair of annealed oligonucleotides were cloned into the single guide RNA clonning site. Two different sgRNAs for human B2M gene were selected Oligonucleotides for clone 1 were: B2M_sg1f CACCGACTCACGCTGGATAGCCTCC and B2M_sg1r AAACGGAGGCTATCCAGCGTGAGTC. Oligonucleotides for clone 2 were: B2M_sg2f CACC. GCAGTAAGTCAACTTCAATGT and B2M_sg2r AAACACATTGAAGTTGACTTACTG. Lentiviral plasmids containing B2M sgRNA were obtained by cloning the hybridized oligonucleotide pairs B2M_sg1f/B2M_sg1r and B2M_sg2f/B2M_sg2r to obtain plasmids pLentiCRISPRv2-B2M1 and LentiCRISPRv2-B2M2 respectively. HeLa cells were transduced with lentiviral preparations of LentiCRISPRv2-B2M1 or LentiCRISPRv2-B2M2 and puromycin resistant clones were selected. After western blotting analysis, clones that did not express B2M were selected, expanded and verified by sequencing the genomic target region.

### VV entry and Blocking assays

For VV entry assays, luciferase activity driven from a VV early promoter at 3 h.p.i. was measured. Cells were plated in MW96 wells and incubated with VV e.luc (m.o.i. = 0.8) for 1 h for adsorption. Cells were washed with PBS and incubated for two additional hours at 37°C with media free of phenol red. Cells were lysed using ONE-Glo EX Luciferase Assay System (Promega) and luciferase activity was measured using EnSight Multimode Plate Reader (PerkinElmer). Blocking assays were performed as follows: for antibody blocking assay HeLa cells were incubated with two different concentrations of anti-B2M antibody (5 and 15 µg/mL) or anti-caveolin antibody (15 µg/mL) -used as negative control- 1 h at room temperature. Subsequently, the cells were infected with V-e.luc (m.o.i.= 0.8) for 1 h and then washed with fresh medium. After 3 h.p.i., cells were lysed and luciferase activity was measured. For B2M blocking assay, V-e.luc inoculum was incubated with growing concentrations of soluble BSA (control) or B2M (0, 5, 15, 30 and 50 µg/mL) 1 h at room temperature. Then, HeLa cells were incubated with the pretreated inocula for 1h and finally washed with fresh medium. At 3 h.p.i. cell extracts were prepared and subsequently luciferase activity was measured.

### Immunofluorescence microscopy

Cells were seeded in round glass coverslips, washed with PBS, fixed with ice-cold 4 % paraformaldehyde for 12 min and permeabilized in PBS containing 0.1% Triton X-100 for 15 min at room-temperature. Cells were treated with PBS containing 0.1 M glycine for 5 min and incubated with primary antibodies in PBS-20 % FBS for 30 min, followed by incubation with secondary antibodies diluted 1:300 in PBS-20 % FBS.

To detect virus particles by antibody labeling on the cell surface, immunofluorescence was carried out on non-permeabilized cells treated as follows: cells were seeded in coverslips, washed and incubated at 12 °C for 1 h with primary antibodies in PBS-20 % FBS followed by incubation at 12 °C for 1 h with secondary antibodies in PBS-20 % FBS. Subsequently, cells were fixed by ice-cold 4 % paraformaldehyde for 12 min and treated with PBS containing 0.1 M glycine for 5 min. Quantitation was derived from counting virus particles in 20-30 cells per sample.

Antibodies used were: Rabbit monoclonal anti-beta 2 Microglobulin EP2978Y antibody (abcam, ref ab75853) diluted 1:300; Mouse monoclonal anti-Integrin beta 1 [12G10] antibody (abcam, ref ab30394) diluted 1:70; Mouse monoclonal anti-VAC (WR) A27L (Beiresources, ref: NR-569) diluted 1:250; anti-mouse IgG-Alexa Fluor 488, anti-mouse IgG-Alexa Fluor 594, anti-rabbit IgG-Alexa Fluor 488 antibodies (Invitrogen) diluted 1:300. Mouse monoclonal anti-Flag 1:500 (Sigma F1804).

For DNA staining, cells in glass coverslips were incubated with 2 mg/mL bisbenzimide (Hoechst dye. Sigma) for 30 min. Cell surface staining was performed by incubation with 0.02 mg/mL wheat germ agglutinin (WGA)- Alexa488 (ThermoFisher Scientific) incubation at concentration for 30 min at 12°C.

For internalization assay, cells were seeded in coverslips, and VV A4-cherry (m.o.i. = 4) was incubated 1 h at 4°C to allow viral adsorption. Cells were then incubated at 37°C for different times in a 5 % CO_2_ incubator to allow internalization. Cells were washed three times with PBS and incubated with antibodies following the non-permeabilization method described above.

Binding assays were based on protocols described elsewhere (6, 20, 82, 83). Briefly, cells grown in coverslips were infected with VV A4-cherry for 1 h at 4 °C (m.o.i. = 30), washed with ice-cold PBS and fixated with paraformaldehyde 4 % for 12 min at 4 °C. Cells were treated with PBS containing 0.1 M glycine for 5 min and incubated 30 min with 0.02 mg/mL Wheat Germ Agglutinin-Alexa488 diluted in PBS-20 % FBS. Cells were washed with PBS, mounted with FluorSave reagent (Millipore) and observed by fluorescence microscopy.

### Western blotting

Whole cell lysates were prepared in denaturant buffer (80 mM Tris-HCl, pH 6.8, 2 % sodium dodecyl sulfate [SDS], 10 % glycerol, 0.01 % bromophenol blue solution and 0.71 M 2-mercaptoethanol). After SDS-polyacrylamide gel electrophoresis (PAGE), proteins were transferred to PVDF membranes and incubated 1 h at RT with primary antibody in PBS containing 0.05 % Tween-20 and 1 % nonfat dry milk. Primary antibodies were Rabbit monoclonal anti-beta 2 Microglobulin EP2978Y antibody (abcam, ref ab75853) diluted 1:100; Rabbit monoclonal anti-Axl C89E7 antibody (Cell Signaling Technology, ref 8661) diluted 1:1000; Rabbit polyclonal anti-beta Actin antibody (GeneTex, ref GTX109639) diluted 1:1000 and Mouse monoclonal anti-Flag 1:1000 (Sigma F1804). After extensive washing with PBS-0.05 % Tween-20, membranes were incubated with HRP-conjugated secondary antibodies diluted 1:3000. Secondary antibodies were polyclonal goat anti-rabbit IgG (Dako P0448), and anti-mouse polyvalent immunoglobulins (Sigma, ref: A0412). After removal of unbound antibody, membranes were incubated for 1 min with a 1:1 mix of solution A (2.5 mM luminol [Sigma], 0.4 mM p-coumaric acid [Sigma], 100 mM Tris-HCl, pH 8.5) and solution B (0.018 % H_2_O_2_, 100 mM Tris-HCl, pH 8.5) to finally record the luminiscence using a Molecular Imager Chemi Doc-XRS (Bio-Rad).

## AKNOWLEDGEMENTS

We are grateful to Dr. Wen Chang for plasmid pm-cherry-A4L and Jerson Garita-Cambronero for assistance with computer analysis. Our gratitude to Dr. Feng Zhang for making the CRISPR/Cas human pooled libraries available through Addgene. We want to thank Marco Y. Hein for critical reading of the manuscript. A.M. was recipient of a predoctoral contract from Subprograma Estatal de Formación, Programa Estatal de Promoción del Talento y su Empleabilidad en I+D+I, Spain.

## Supporting information

**S1 Fig. Screening experiment example**. A representative hit analysis for one of the 27 screen experiments analyzed with MaGeCK. For each gene, the LFC values from infected and non-infected cultures are represented. Gene hits with FDR < 0.05 that also show LFC > 0 in infected and LFC < 0 in non-infected are labeled with gene name tags. Best hits are those that have the greatest LFC in infected control and the lowest LFC in control experiments (non infected). Note that genes with non infected LFC > 1 could be positive hits as well.

**S2 Fig. VV / B2M colocalization rates.** HeLa cells were infected with VV A4-cherry (red) for 1 h at 4 °C and unbound virus was removed by washing (m.o.i. = 5). Non-permeabilized cells were then incubated for different times at 37 °C and stained with anti-B2M to analyze colocalization.

**S3 Fig. Blocking assay for VV entry by soluble B2M.** V-e.luc virus was incubated with increasing concentrations of soluble BSA (control) or B2M (0, 5, 15, 30 and 50 µg/mL) protein for 1 h at room temperature. Then, HeLa cells were incubated for 1 h with the pre-treated inoculum. At 3 h.p.i. luciferase activity was determined as a measure of viral entry and early gene expression. No significant differences were found. p-values: **** < 0.0001, *** < 0.001, ** < 0.01, * < 0.05, ns > 0.05.

**S4 Fig. Blocking assay for VV entry with anti-B2M antibody.** HeLa cells were incubated with two different concentrations of anti-B2M antibody (5 and 15 µg/mL) for 1 h at room temperature. Anti-caveolin antibody (15 µg/mL) was used as negative control. After antibody treatment, HeLa cells were infected with V-e.luc (m.o.i. 0.8), and eventually 3 h.p.i. luciferase activity was determined as a measure of viral entry and early gene expression. No significant differences were found. ns, not significant (p> 0.05).

**S1 Table. Screen_conditions.** Different screen experiments were performed. Experiments are summarized where which mutant, m.o.i. and number of reinfections is showed.

**S2 Table. Screen Results.** Gene summary files obtained from MaGeCK test showing the FDR and LFC for each gene in each experiment.

**S3 Table. ScreenBEAMresults.** Summary of ScreenBEAM results showing good guides, B-score, z-score, p-value, FDR and standard deviations for each gene, where x is for control experiment and y for infected experiment. Top 20 genes were selected as hits.

**S4 Table. Reactome_Pathway analysis**. Results from Reactome Pathway analysis. The most enriched pathways are shown in descendent order. Statistical data is shown (i.e. pvalue, p.adjust), as well as the genes ID that belong to each enriched pathway.

